# Respiration is essential for aerobic growth of *Zymomonas mobilis* ZM4

**DOI:** 10.1101/2023.03.02.530925

**Authors:** Magdalena M. Felczak, Michaela A. TerAvest

## Abstract

*Zymomonas mobilis* is an alpha-proteobacterium that is a promising platform for industrial scale production of biofuels or valuable products due to its efficient ethanol fermentation and low biomass generation. *Z. mobilis* has also intriguing physiology, sometimes difficult to explain by the rules and strategies commonly observed in other bacteria. One of the most mysterious features of *Z. mobilis* is its growth in oxic conditions. *Z. mobilis* is an aerotolerant bacterium that encodes a complete respiratory electron transport chain but the benefit of respiration for growth in oxic conditions has never been confirmed, despite decades of research. Quite the opposite, growth and ethanol production of WT *Z. mobilis* is poor in oxic conditions indicating that it does not benefit from oxidative phosphorylation. Additionally, in previous studies, aerobic growth improved significantly when respiratory genes were disrupted (*ndh*) or acquired point mutations (*cydA, cydB)* even if respiration was significantly reduced by these changes. Here, we obtained clean deletions of respiratory genes *ndh* and *cydAB*, individually and in combination, and showed, for the first time, that deletion of *cydAB* completely inhibited O_2_ respiration and dramatically reduced growth in oxic conditions. Both respiration and aerobic growth were restored by expressing a heterologous, water-forming NADH oxidase, *noxE*. This result shows that the main role of the electron transport chain in *Z. mobilis* is reducing the toxicity of molecular oxygen, helping to explain why it is beneficial for *Z. mobilis* to use electron transport chain complexes that contribute little to oxidative phosphorylation.

**Importance:** A key to producing next generation biofuels is to engineer microbes that efficiently convert non-food materials into drop-in fuels and to engineer microbes effectively we must understand their metabolism thoroughly. *Zymomonas mobilis* is a bacterium that is a promising candidate biofuel producer but its metabolism remains poorly understood, especially its metabolism when exposed to oxygen. Although *Z. mobilis* respires with oxygen, its aerobic growth is poor and disruption of genes related to respiration counterintuitively improves aerobic growth. This unusual result has sparked decades of research and debate regarding the function of respiration in *Z. mobilis*. Here, we used a new set of mutants to determine that respiration is essential for aerobic growth and likely protects the cells from oxidative damage caused by molecular oxygen. These results indicate that respiration has a non-canonical function in *Z. mobilis* and expand our understanding of the role of respiration in metabolism and oxidative stress.

## Introduction

*Zymomonas mobilis* is a facultative anaerobe of interest for biofuel production from sugars. There are also several unusual characteristics of *Z. mobilis* that make it an interesting model for testing assumptions about bacterial physiology and metabolism. For example, unlike other preferentially anaerobic organisms, *Z. mobilis* uses the Entner-Doudoroff pathway for glycolysis (1 ATP per glucose) rather than the Embden-Meyerhof-Parnas pathway (2 ATP per glucose) (1). Additionally, attempts to genetically modify *Z. mobilis* have yielded evidence that this organism carries multiple copies of its genome, with some estimates as high as 50 copies per cell (2, 3). The aerobic metabolism of *Z. mobilis* is also unusual and has been the subject of debate. Although *Z. mobilis* has a documented respiratory electron transport chain (ETC), it grows more quickly and to a higher density under anoxic conditions than under oxic conditions (4, 5). The poor aerobic growth of *Z. mobilis* is surprising because respiration generates more ATP per glucose than fermentation and therefore, facultative aerobes typically have a higher growth yield (mg biomass dry weight/ mg glucose) in the presence of oxygen. The negative effect of oxygen on *Z. mobilis* has been attributed to accumulation of the toxic metabolic intermediate acetaldehyde (6, 7). While some of the acetaldehyde is converted to acetate by an acetaldehyde dehydrogenase, residual acetaldehyde is often observed in aerobic cultures of *Z. mobilis* (8).

The respiratory chain of *Z. mobilis* also has a low predicted coupling efficiency, i.e., few ATP are generated per O_2_ consumed, possibly leading to a smaller than usual benefit for using the ETC. Based on the genome sequence, *Z. mobilis* is predicted to have 4 dehydrogenases that donate electrons to the quinone pool: NAD(P)H dehydrogenase (Ndh, ZMO1113), quinone-linked lactate dehydrogenase (Ldh, ZMO0256), glucose dehydrogenase (Gdh, ZMO0072), and succinate dehydrogenase (Sdh, ZMO0568-0569). A *bd*-type terminal oxidase (CydAB, ZMO1571-1572) and a *bc1* complex (quinol:cytochrome *c* oxidoreductase, ZMO0956-0958) are also encoded in the genome, but there is no known cytochrome *c* oxidase to complete the *bc*_*1*_ pathway. Because none of the predicted ETC complexes are proton pumping, proton translocation only occurs through the oxidation and reduction of respiratory quinones at active sites on opposite sides of the membrane (**Figure 1**). Compared with ETC configurations in other organisms that may translocate as many as 10 H^+^/2 e^-^, the coupling efficiency of 2 H^+^ /2 e^-^ in *Z. mobilis* is very low and may not appreciably contribute to ATP synthesis.

**Figure 1.**
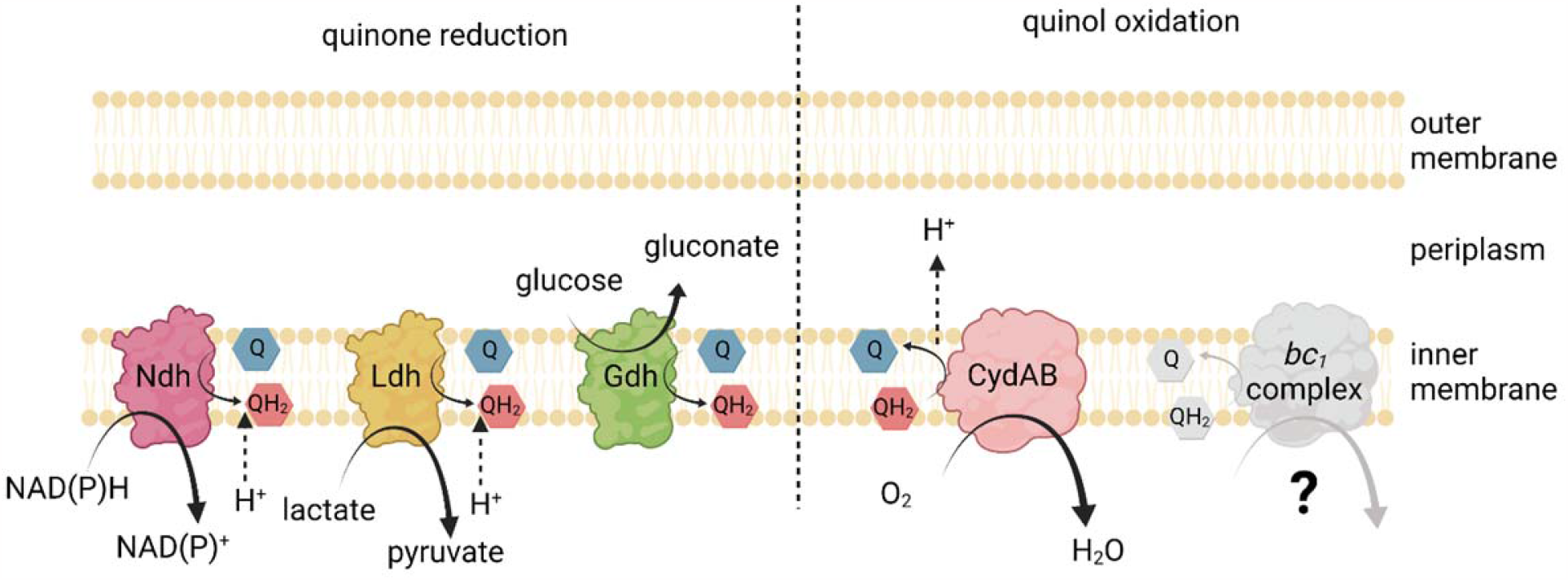
Proposed structure of the electron transport chain in *Z. mobilis*. This schematic depicts electron transport complexes encoded in *Z. mobilis*. Lactate dehydrogenase (Ldh), NADH dehydrogenase (Ndh), glucose dehydrogenase (Gdh), *bd*-type quinol oxidase (CydAB), and quinol-cytochrome *c* oxidoreductase (*bc*_*1*_ complex). Not shown: succinate dehydrogenase (Sdh). Created with BioRender.com.

Multiple research groups have previously shown that disruption of the gene encoding the NADH dehydrogenase in *Z. mobilis* (*ndh*) improves aerobic growth (9–12). The improved growth of respiratory mutants has raised the question of why ETC genes are conserved in the genome, since they appear to have no effect under anoxic conditions and to be deleterious under oxic conditions. Jones-Burrage et al. (9) found evidence that while respiratory chain activity does not enhance growth, it may contribute to survival in stationary phase in minimal media. One complicating factor for studies that disrupt Ndh is that there are other entry points into the electron transport chain, including Gdh, Ldh, and Sdh, indicating that disrupting Ndh does not completely block the ETC (13). Further, in the absence of Ndh, the respiratory (quinone-linked) lactate dehydrogenase and the cytosolic (NAD^+^-linked) lactate dehydrogenase (ZMO1257) may form a bypass whereby NADH can be oxidized and electrons enter the respiratory chain in the absence of functional Ndh (14).

There are also unusual aspects of the latter half of the ETC. The *Z. mobilis* genome encodes a *bd* oxidase and a *bc*_*1*_ complex but no cytochrome *c* oxidase that would be predicted to terminate the *bc*_*1*_ branch. Sootsuwan et al. proposed that a peroxidase could terminate this branch using either hydrogen peroxide or O_2_ as the electron acceptor (15). A later study determined that while the peroxidase PerC did contribute to hydrogen peroxide tolerance, it did not accept electrons from the *bc*_*1*_ complex or contribute significantly to the oxygen consumption rate of *Z. mobilis* (16). Strazdina et al. previously disrupted genes in the *bc*_*1*_ complex (*cytB*) and *bd* complex (*cydB*) by a chloramphenicol resistance marker insertion (17). These mutants had very low oxygen consumption rates initially but recovered WT-level oxygen consumption rates after 11 to 12 hours of aerobic growth. This could have been due to changes in regulation or instability of the mutation. As observed by others, disruption of the *bc*_*1*_ complex had little effect on growth.

Overall, previous studies of the ETC in *Z. mobilis* have identified active complexes but have not determined whether they contribute to proton motive force (PMF) generation and oxidative phosphorylation. In this study, we deleted *ndh* and *cydAB* individually and in combination and measured growth, glucose metabolism, and oxygen consumption rates in the mutant strains. In contrast to previous studies, we found that ETC flux is essential to *Z. mobilis* in oxic conditions. We complemented an ETC-deficient mutant with a water-forming NADH oxidase (NoxE), which oxidizes NADH and consumes O_2_ but does not contribute to proton-motive force to drive oxidative phosphorylation. Results with NoxE indicated that oxygen removal from the environment is a key role of the ETC but not the sole benefit it provides to *Z. mobilis*.

## Results

### Construction of mutant strains

To investigate the role of the electron transport chain in *Z. mobilis*, we constructed deletion mutants of the genes encoding the NADH dehydrogenase (*ndh*, ZMO1113) and the *bd* quinol oxidase (*cydAB*, ZMO1571-72) using a homologous recombination method (18). Briefly, regions upstream and downstream of the target gene(s) are cloned into a vector backbone that also contains a constitutively expressed *gfp*, antibiotic resistance marker, and an origin that cannot replicate in *Z. mobilis*. The resulting plasmid is transferred to *Z. mobilis*, and antibiotic resistant colonies are obtained when the plasmid integrates into the genome by homologous recombination. A second recombination event removes the plasmid from the genome, leaving either the WT or deletion sequence. All growth steps of the procedure were performed in anoxic conditions to avoid selection against the mutant strains (see Materials and Methods). Deletion of *ndh* by this method was straightforward, but deletion of *cydAB* was more challenging. Colonies that had undergone the second recombination event were identified by the loss of GFP fluorescence. These were screened for deletion by PCR amplification of the region of interest. For the *cydAB* deletion, we observed that most colonies that produced the PCR amplicon consistent with deletion also produced an amplicon consistent with the WT sequence. Of 39 colonies screened by PCR, 6 showed the deletion band and only one of these also lacked the WT band (**Figure S1**, isolant 5). For comparison, 8 colonies of 11 screened for *ndh* knockout showed the deletion band, and none of these contained the wild type band with the deletion band. The *ΔcydAB* strain was also checked by Illumina re-sequencing and we found no reads aligned in the deleted region (**Figure S2**). We also generated a Δ*cydAB*Δ*ndh* deletion strain by deleting *ndh* from the Δ*cydAB* strain.

### Growth and metabolism of mutant strains

As expected, the growth of both Δ*ndh* and Δ*cydAB* strains was indistinguishable from WT ZM4 in anoxic conditions (**Figure S3**). We observed that deletion of *ndh* improved growth in oxic rich medium (**Figure 2A**). In contrast, deletion of *cydAB* dramatically reduced growth in the same conditions. Complementation of the deletions with the corresponding genes expressed from a plasmid returned each strain to the WT phenotype, decreasing growth of Δ*ndh* and increasing growth of Δ*cydAB*. Note that to facilitate comparisons, the non-complemented strains in Figure 2 contain the vector, pRL814, an inducible plasmid carrying *gfp* that was used to construct both complementing plasmids. Here, we will refer to pRL814 as ‘vector’. This plasmid does not change the growth of WT or knockout strains when uninduced (data not shown). We observed the most complete growth complementation without IPTG induction and increasing IPTG concentration made complementation less effective (Figure S4). We have previously observed that expression from pRL814 is leaky, so some expression of the gene of interest occurs even when inducer is not added (19).

**Figure 2.**
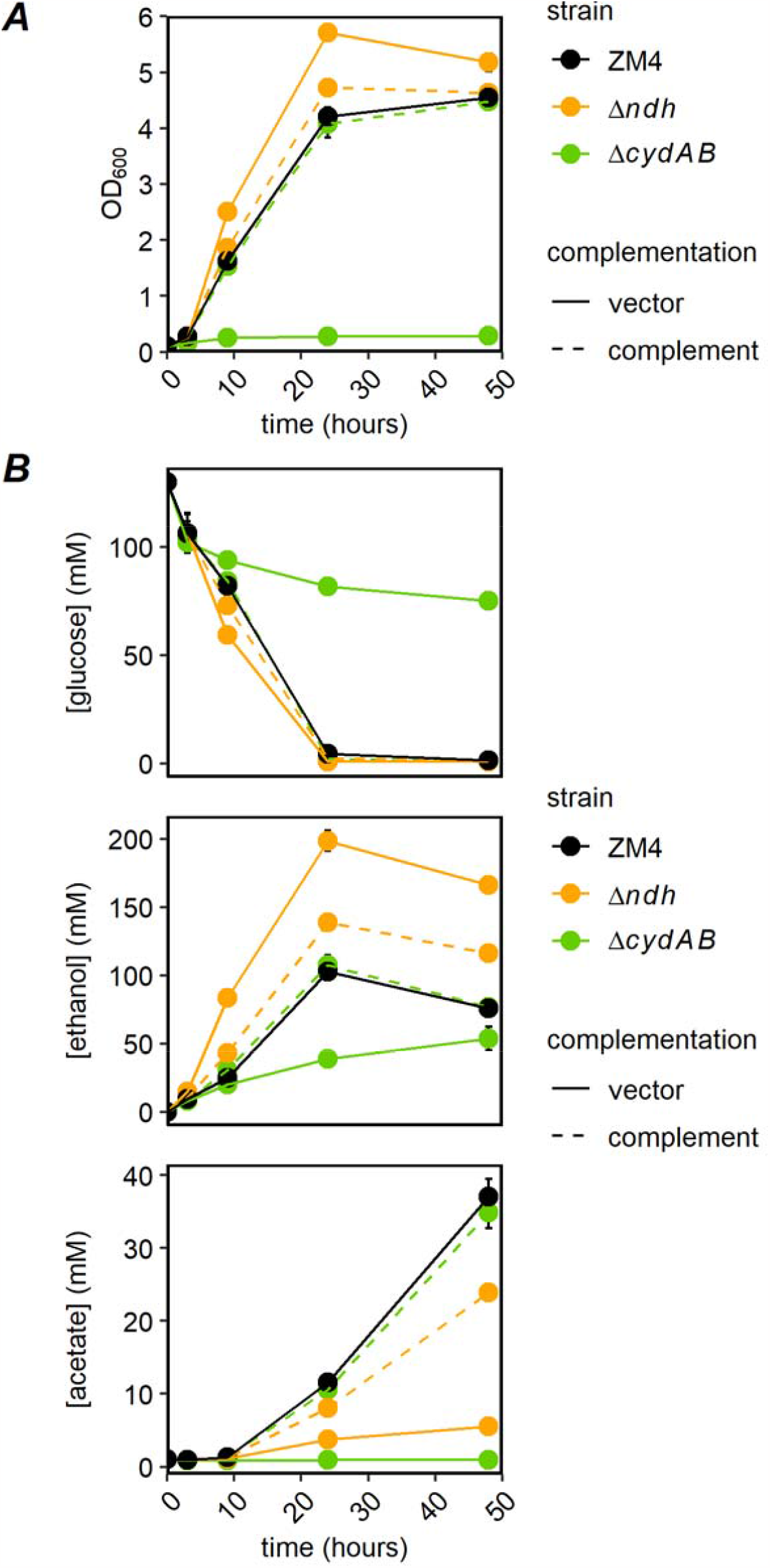
(A) Growth and (B) HPLC analysis of ZM4 and Δ*cydAB* and Δ*ndh* in oxic rich medium, with and without complementation of deleted genes. ZM4 and Δ*cydAB* and Δ*ndh* bearing the vector or respective complementing plasmids were grown in rich medium (ZRMG) with spectinomycin in an anaerobic chamber overnight. In the morning, cultures were diluted to OD_600_ = 0.1 in 5 ml of fresh aerobic ZRMG with spectinomycin and grown in glass tubes covered with loose caps with shaking at 30°C at ambient oxygen pressure for 48 hours. At times indicated, OD_600_ was measured and samples were centrifuged. Supernatants were analyzed by HPLC as described in “Materials and Methods”. Metabolite concentrations were calculated from standard curves. Points are the average of three biological replicates and error bars are standard errors. Complement: pRL*ndh* or pRL*cydAB*.

Consistent with the improved growth, deletion of *ndh* increased glucose consumption and ethanol production and reduced acetate production compared with WT (**Figure 2B**). The Δ*cydAB* strain showed low glucose consumption and ethanol and acetate production, consistent with the poor growth. In oxic conditions, both mutant strains produced ethanol more efficiently than WT, with WT producing 29% of the theoretical yield, Δ*ndh* producing 65%, and Δ*cydAB* producing 49%. Thus, although the Δ*cydAB* strain produced less ethanol than WT, it converted glucose to ethanol more efficiently. Again, both strains were returned to the WT phenotype (i.e., low efficiency ethanol production) by expressing the deleted gene from a plasmid.

We measured the oxygen consumption rate of each strain using microplates that contained optical sensors integrated into the bottom of the wells to facilitate high throughput monitoring of oxygen consumption (Oxoplates). Oxygen partial pressure (pO_2_) is measured as changes in fluorescence of the optical sensor, which is oxygen dependent. Cultures grown anaerobically overnight were diluted in fresh aerobic medium to an OD_600_ appropriate for measuring pO_2_ over time. For example, fast respiring strains, like wild type ZM4, were diluted to OD_600_ = 0.2, while deletion mutants were diluted to OD_600_=1.0. Cultures were aerated for 30 minutes by shaking in a plate reader; this time should be sufficient to induce any ETC genes that are upregulated by oxygen exposure (20). Next, measurements were taken without shaking for 60 minutes. The final rate of oxygen uptake was normalized to OD_600_=1. We found that ZM4 consumed oxygen at a rate of 0.941±0.017 mg/L/min, while Δ*ndh* consumed oxygen at 13% of the WT rate and Δ*cydAB* consumed oxygen at 0.6% of the WT rate (**Figure 3**). Complementing Δ*ndh* with *ndh* returned the oxygen consumption rate to 60% of the WT rate and complementation of Δ*cydAB* with *cydAB* returned oxygen consumption to 106% of the WT rate (representative dissolved oxygen profiles in Figure S5).

**Figure 3.**
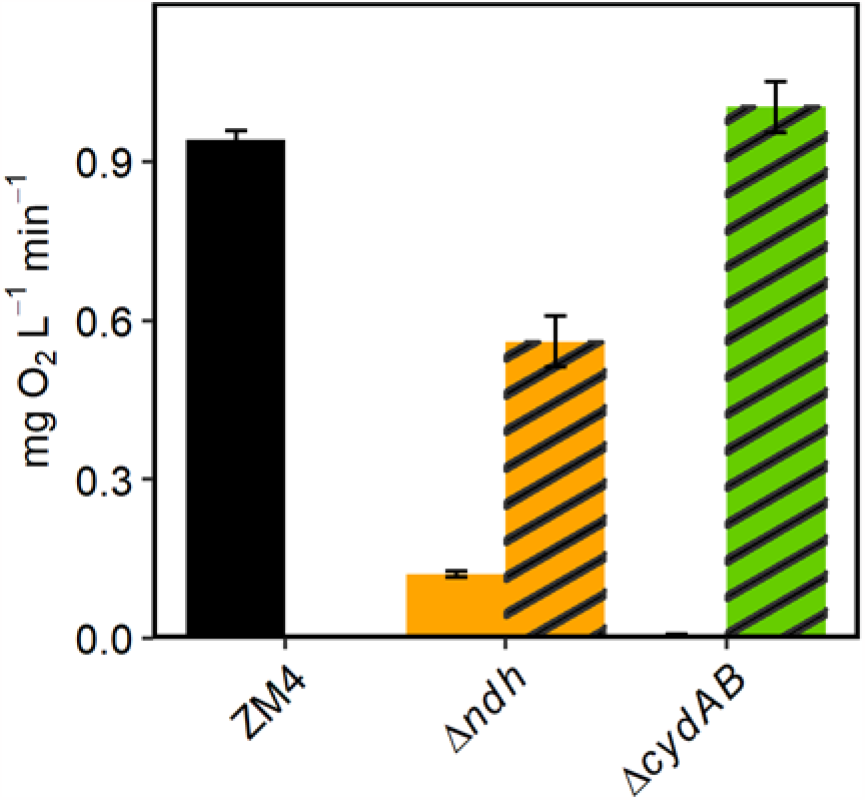
Oxygen consumption by *Z. mobilis* ZM4 and Δ*ndh* and Δ*cydAB* mutants with and without complementation. Strains were grown in rich medium (ZRMG) with spectinomycin at 30°C in an anaerobic chamber, overnight. Overnight cultures were diluted in fresh ZRMG with spectinomycin to an OD_600_ appropriate for oxygen measurement (see text); ZM4/vector, and Δ*ndh* and Δ*cydAB* strains with complementing plasmids were diluted to OD_600_ = 0.2, and Δ*ndh* and Δ*cydAB* bearing the vector to OD_600_ = 1.0. 200 µl from each dilution was loaded onto an Oxoplate in triplicate. Plates were incubated with shaking at 30°C in a plate reader for 30 minutes. After this time, shaking was stopped and fluorescence was measured every three minutes for 60 minutes. Oxygen partial pressure (pO_2_) as “% air saturation” was calculated from two-point calibration of the Oxoplate, as described in “Materials and Methods”. A conversion factor of 0.091 was used to convert “% air saturation” to “mg O_2_/L”. Oxygen uptake rate in mg/L/min, was calculated from the linear part of the curve and normalized to OD_600_=1.0. Bar graphs are average of three biological and three technical replicates and error bars are standard errors. Striped bars, strains with complementing plasmids. Note that there is only one column for ZM4 with no plasmid. The bar for Δ*cydAB* is too small to be seen on this scale.

### Analysis of a Δ*cydAB*Δ*ndh* double mutant

One possible explanation for the difference in growth phenotypes between the Δ*ndh* and Δ*cydAB* strains, despite both being part of the same metabolic pathway, is that these two mutations impact the redox state of the quinone pool in different ways. Removal of Ndh would make the quinone pool more oxidized while removal of CydAB would make it more reduced. Overreduction of the quinone pool may be toxic, possibly leading to the negative effect of the Δ*cydAB* mutation. To test this, we generated a double mutant lacking both *ndh* and *cydAB*. If quinone overreduction was responsible for poor growth of Δ*cydAB*, removing *ndh* should alleviate or decrease this effect. As expected, the oxygen uptake rate of the double mutant was negligible, as in the Δ*cydAB* strain. Complementing with pRL*ndh*, did not improve the oxygen consumption and complementing with pRL*cydAB* increased it to a level similar to the Δ*ndh* strain (**Figure 4**, representative profiles in Figure S6). As in previous figures, strains containing the vector were used as the negative control for complementation.

**Figure 4.**
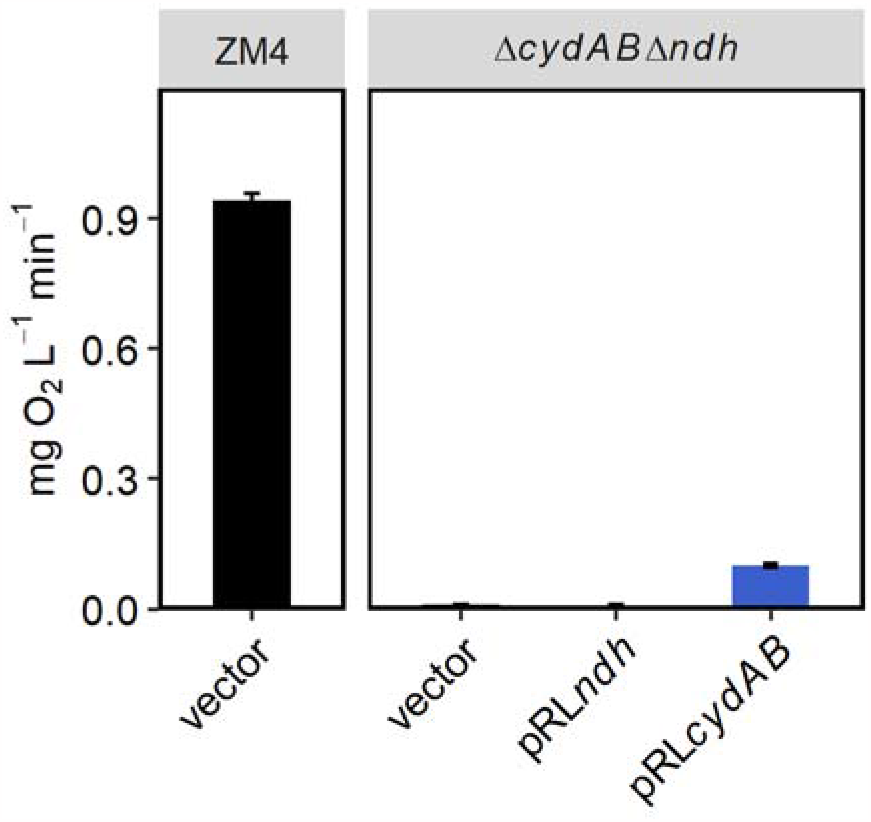
Oxygen consumption by Δ*cydAB*Δ*ndh* with complementing plasmids. Strains were grown as described in Figure 3. ZM4/vector was diluted to OD_600_ = 0.2 while the double mutant strains bearing the vector or a complementing plasmid were diluted to OD_600_ = 1.0. Oxygen partial pressure was measured and O_2_ uptake rate was calculated as described in “Materials and Methods” and in Figure 3. O_2_ uptake rate in mg/L/min was normalized to OD_600_=1.0. Bar graphs are average of three biological and three technical replicates and error bars are standard errors.

We observed that the double mutant had the same phenotype as Δ*cydAB*, growing very poorly in oxic conditions (**Figure 5A**). Similar to the observation for oxygen consumption, complementation with *cydAB* rescued growth, while complementation with *ndh* had no effect on growth. Glucose consumption, ethanol production, and acetate production for the double mutant and complemented strains were all consistent with the growth, essentially as in the Δ*cydAB* strain (**Figure 5B**). These results indicate that overreduction of the quinone pool is not a likely reason for the difference in phenotype between Δ*ndh* and Δ*cydAB*. It also confirms that Ndh and CydAB are operating in the same metabolic pathway (ETC).

**Figure 5.**
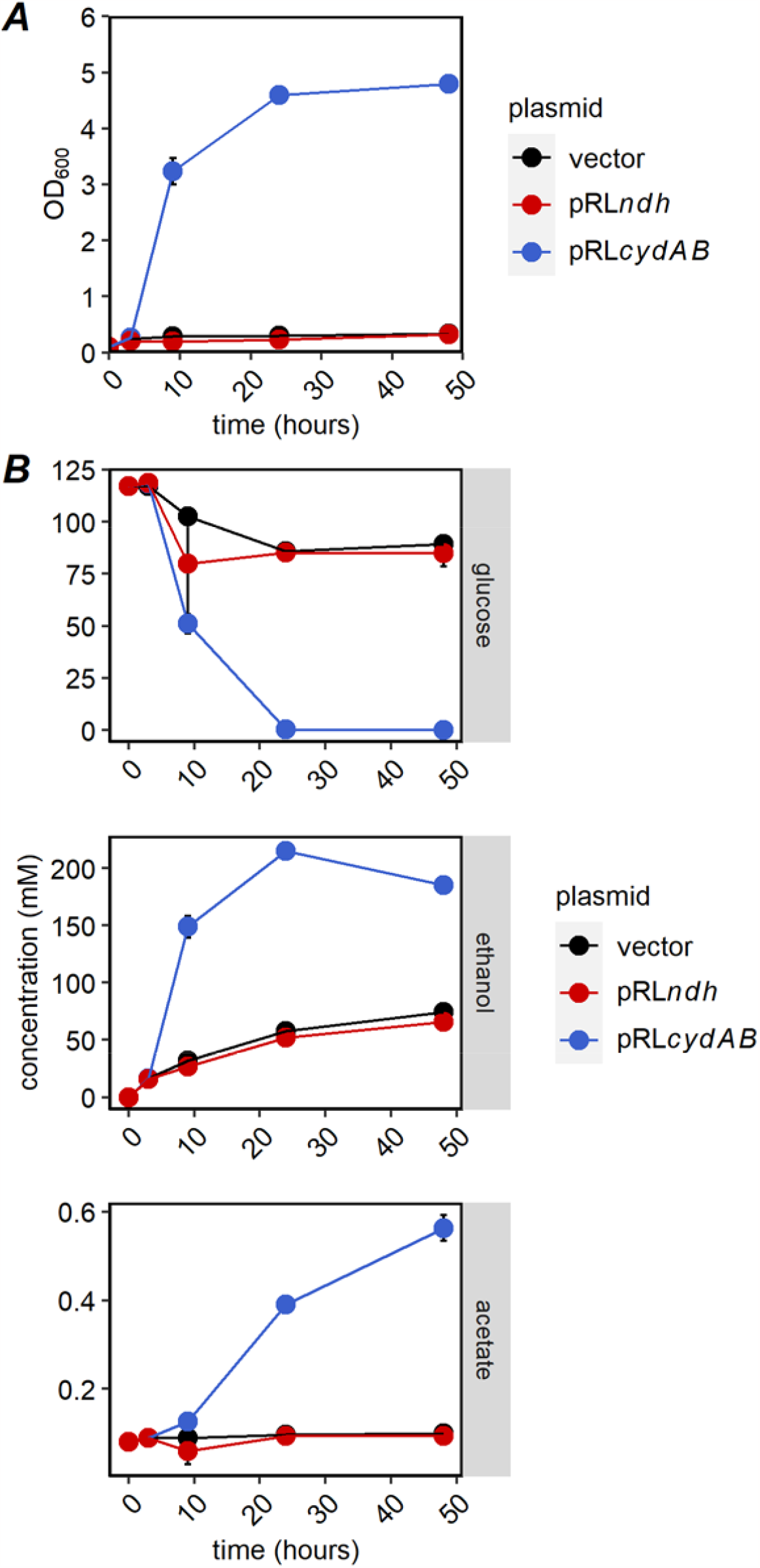
(A) Growth and (B) HPLC analysis of Δ*cydAB*Δ*ndh* with complementation. Δ*cydAB*Δ*ndh* strain bearing the vector or one of the complementing plasmids were grown in ZRMG with spectinomycin as described in Figure 2. OD_600_ was measured at times indicated to monitor growth and supernatants were analyzed by HPLC as described in Figure 2. Points are average of three biological replicates and error bars are standard errors.

### Complementation with a water-forming NADH oxidase

CydAB may be important to *Z. mobilis* for different reasons, including reoxidation of quinols in the inner membrane, removal of O_2_ from the environment for protection from reactive oxygen species, or contribution to PMF generation for ATP synthesis or other processes. To determine whether oxygen consumption was the key role of CydAB, we recombinantly expressed NoxE, a water-forming NADH oxidase that oxidizes NADH and reduces O_2_ to H_2_O in the cytoplasm. NoxE consumes oxygen without contributing to PMF formation. When *noxE* was expressed in *Z. mobilis* Δ*cydAB*, we observed IPTG-dependent oxygen consumption, indicating that functional NoxE was produced (**Figure 6**). Maximal oxygen consumption was observed with induction with 100 µM IPTG. At this induction level, oxygen consumption of the Δ*cydAB* strain producing NoxE was 56% of the WT rate. We also measured growth at a range of IPTG concentrations and observed that NoxE expression moderately improved growth of Δ*cydAB* (**Figure 7**).

**Figure 6.**
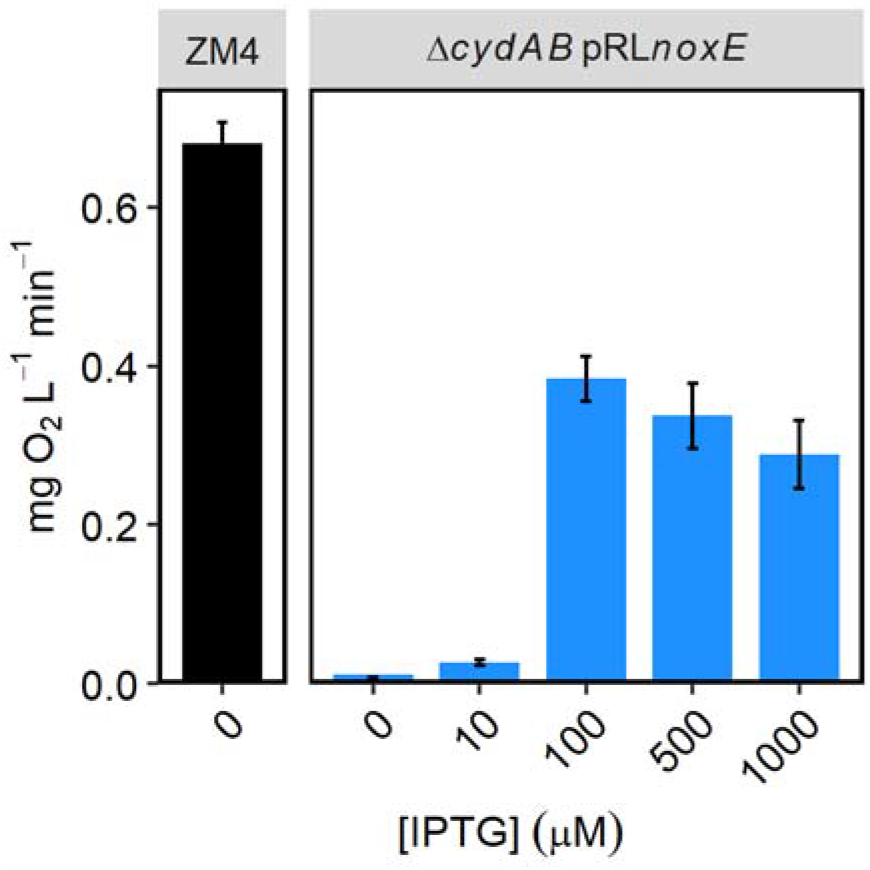
Oxygen consumption with heterologous expression of *Lactobacillus brevis noxE*. Δ*cydAB*/pRL*noxE* strain was grown in rich defined medium (ZRDM) containing spectinomycin and indicated concentrations of IPTG, at 30°C in an anaerobic chamber, overnight. ZM4/vector was grown without IPTG. Δ*cydAB*/pRL*noxE* cultures were diluted to OD_600_ of 0.2, 0.5 and 1.0 in ZRDM containing spectinomycin and the same level of IPTG as the overnight culture. ZM4/vector was diluted to OD_600_=0.2 in ZRDM without IPTG. 200 µl of each dilution was loaded on Oxoplate, in triplicate. Oxygen partial pressure (pO2) was measured as described in Materials and Methods and in Figure 3. pO_2_ from measurable dilutions were normalized to OD_600_=1.0. Bar graphs show average oxygen uptake rate after normalization. N ≥6 for Δ*cydAB*/pRL*noxE* and n=3 for ZM4/vector.

**Figure 7.**
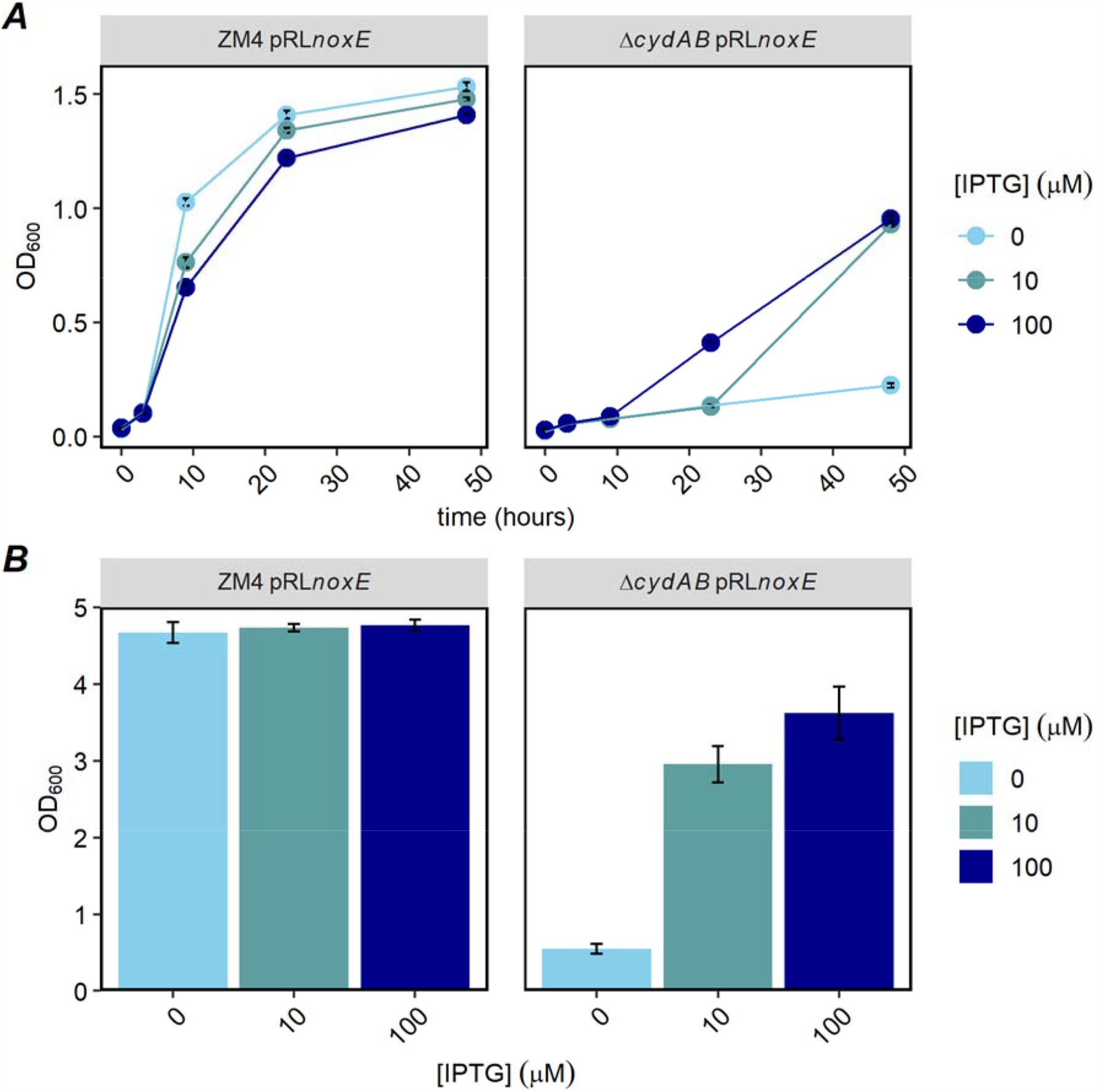
Growth of Δ*cydAB* with heterologous expression of *Lactobacillus brevis noxE*. ZM4 and Δ*cydAB* **s**trains, bearing the vector or pRL*noxE* were grown in rich defined medium (ZRDM) with spectinomycin at 30°C in an anaerobic chamber, overnight. Cultures were diluted in fresh ZRDM, supplemented with spectinomycin, to OD_600_=0.1. IPTG was added to final concentrations 0, 10 and 100 µM. 200 µl from each culture at indicated IPTG level was loaded on 96-well clear microplate, in triplicate. Cells were grown on a plate shaker in 30°C at ambient oxygen pressure for 48 hours. Growth was monitored by measuring absorbance at 600 nm in a plate reader. After 48 hours, cultures from three wells were combined and final OD_600_ was measured by a spectrophotometer with a 1-cm pathlength. **A**: growth curve from a representative experiment measured by plate reader; average of three technical replicates. **B**: OD_600_ after 48 hours of incubation; average from three independent plates, with three technical replicates each. Error bars are standard error. In **A** and **B** only strains bearing pRL*noxE* are shown because ZM4 and ZM4/vector grow the same, and Δ*cydAB*/vector grows as uninduced Δ*cydAB*/pRLnoxE or less when induced.

## Discussion

As previously observed by other groups, we found that deletion of *ndh* from the *Z. mobilis* genome resulted in enhanced growth in oxic conditions. However, when we deleted *cydAB* from the genome, we observed a severe growth defect in oxic conditions. These results suggest that contrary to the previous hypotheses that respiration is harmful to *Z. mobilis*, the electron transport chain is essential to aerobic growth. We speculate that all previous respiratory deficient conditions, whether induced by a mutation or inhibitor, likely still had significant respiratory activity, resulting in the lack of a growth defect in previous studies. In the case of Ndh mutation, Strazdina et al. found that removing Ndh alone is insufficient to block respiratory activity because the NAD^+^-linked and quinone-linked lactate dehydrogenases create a bypass that allows entry of electrons into the ETC (14). Our oxygen consumption results corroborate their findings, indicating that electrons can still pass through the ETC in the absence of Ndh. The respiration deficient mutants isolated by Hayashi et al. included some large deletions or stop codons in *ndh* but only point mutations in *cydA* and *cydB*, indicating that all these mutants likely had residual respiratory activity. Similar to Ndh disruption, cyanide addition to *Z. mobilis* cultures reduces the respiratory rate but does not block the ETC completely and leads to enhanced growth in oxic conditions (6).

Strazdina et al. previously generated a *cydAB* mutant via insertion of a chloramphenicol resistance cassette, but found that the mutant retained respiratory capacity (17). They suggested that an alternative pathway involving the *bc1* complex could compensate for the lack of the *bd* oxidase and thus respiration would still occur if only one of the pathways is removed. Our results show that only the CydAB pathway is active in ZM4, leaving the possible role of *bc1* complex unknown. In Strazdina et al., growth data for the *cydAB* mutant strain were not presented, so it is difficult to compare our strain with theirs, although we suspect that, due to chromosomal polyploidy of Zymomonas (2, 3), they may have had a mixture of the disrupted and WT sequences,. Other groups have also observed that intended gene deletions in *Z. mobilis* actually function as knockdowns rather than knockouts because each cell carries multiple genome copies and in some cases not all copies are disrupted (2). The polyploidy of *Z. mobilis* likely led to the hybrid PCR results we observed during construction of Δ*cydAB*, but PCR indicated that we obtained a colony with all copies deleted (Figure S1). Further, genome sequencing did not detect any reads in the Δ*cydAB* open reading frame, indicating that we generated a true deletion mutant (Figure S2).

Comparing the growth and oxygen consumption rates of WT, Δ*ndh*, and Δ*cydAB*, it appears that the optimal oxygen consumption rate for ZM4 in oxic laboratory growth is somewhere between zero and the WT level. Although the ETC appears to be sub-optimally regulated in laboratory conditions, we found that it is necessary for oxic growth. It remains puzzling why *ndh* has been conserved in the *Z. mobilis* genome when deletion seems to have no effect or improve growth in most conditions. Jones-Burrage et al. found that Ndh contributes to survival during stationary phase in oxic minimal medium, suggesting that there may be an important role under environmental conditions (9). Future work to simultaneously delete multiple respiratory dehydrogenases may shed light on the specific role of Ndh.

Previous work suggested that aerobic respiration is harmful to *Z. mobilis* because of acetaldehyde formation (4) and that *ndh* mutants grow better than WT because of reduced acetaldehyde concentrations. However, we observed that the Δ*cydAB* strain grew very poorly despite producing no acetaldehyde (Figure S7). This indicates that acetaldehyde toxicity alone is insufficient to explain the poor aerobic growth of *Z. mobilis* or the growth enhancement of Δ*ndh* strains. To test whether oxygen toxicity itself is the cause of poor growth in Δ*cydAB*, we expressed a heterologous water-forming NADH oxidase, NoxE. We found that NoxE did improve the final culture density to ∼75% of the WT level, indicating that reducing O_2_ concentration in the culture is one of the key functions of the ETC. Previous multi-omic analysis of oxygen exposure in *Z. mobilis* indicated that iron sulfur cluster enzymes were quickly inactivated by O_2_, leading to major disruptions in central metabolism (20). By expressing the electron transport chain, *Z. mobilis* can quickly remove oxygen from the environment (within minutes) and begin repairing iron sulfur cluster enzymes to return to metabolic homeostasis.

Although our results show that oxygen consumption is likely the main role of the ETC, there was no condition under which NoxE fully complemented growth, suggesting that the ETC may also contribute other benefits, possibly including formation of a proton-motive force across the inner membrane. Although the oxygen consumption rate of the NoxE complemented strains was lower than WT, the oxygen consumption for Δ*ndh* was also lower and this strain grew better than WT. Therefore, we do not think the slower oxygen consumption can fully explain why NoxE did not fully rescue the *cydAB* deletion. We attempted to test whether the ETC contributes to proton-motive force generation by expressing a light-driven proton pump, proteorhodopsin, in Δ*cydAB* but we did not observe complementation (data not shown). We were not able to observe any effect of proteorhodopsin expression in any strain background, except for a general growth defect in all strains at high expression levels, possibly due to protein aggregation or disruption of the membrane. Because we could not confirm that proteorhodopsin was functional in *Z. mobilis*, we cannot conclude that generation of proton-motive force is not a role of the ETC. Further work is necessary to understand the role of the ETC in generating PMF in aerobic growth of *Z. mobilis*.

Although respiration is generally considered a source of ATP via oxidative phosphorylation, our results indicate that energy conservation is not the only or even the major role of the electron transport chain in some organisms. Evidence that oxygen consumption is the main role of the ETC in *Z. mobilis* helps to explain why complexes with low coupling efficiency are used. If ATP synthesis via oxidative phosphorylation is not important, proton translocation is not needed and because the less efficient ETC components tend to be smaller and simpler than their efficient counterparts they incur less metabolic cost to produce. For example, the proton-translocating NADH dehydrogenase in *E. coli* consists of 14 subunits, while the uncoupling version has only one. Therefore, by using a low efficiency electron transport chain, *Z. mobilis* can gain the benefits of oxygen consumption with the minimum metabolic cost. Low efficiency electron transport complexes are found in a wide variety of bacteria and future work will further expand our understanding of a likely wide range of functions they can contribute to beyond oxidative phosphorylation.

## Materials and Methods

### Media and Chemicals

*Escherichia coli* strains were grown in Luria-Bertani media (Miller, Acumedia). *Zymomonas* Rich Medium (ZRMG) contains 1% yeast extract, 2% D-glucose and 15 mM KH_2_PO_4_. *Zymomonas* Rich Defined Medium (ZRDM) contains: 40 mM potassium phosphate buffer pH 6.2, 2% glucose, 0.05% NaCl, 0.1% (NH4)_2_SO_4_, 0.2 g/L MgSO4 × 7H_2_O, 0.025 g/L of Na_2_MoO_4_ x 2H_2_O, 0.01 g/L of CaCl_2_ x 2 H_2_O, 0.0025 g/L FeSO_4_ x 7 H_2_O, 0.001 g/L calcium pantothenate, 1 x Teknova EZ supplement, and 1 x Teknova ACGU solution. 2,6-diamino pimelic acid (DAP) was added to 0.3 mM when indicated. Spectinomycin and chloramphenicol were used at 100 µg/ml or 70 µg/ml, respectively, for *Zymomonas* or at 50 µg/ml or 35 µg/ml, respectively for *E. coli*. IPTG (Sigma) was added as indicated. Restriction enzymes, T4 Ligase, Q5 High Fidelity Polymerase and HiFi DNA assembly master mix were from New England Biolabs. Oligonucleotides and gBlocks were from IDT.

### Bacterial strains and plasmids

*E. coli* Mach 1 competent cells were from ThermoFisher Scientific. *Zymomonas mobilis mobilis* ATCC 31821 (ZM4) was from American Type Culture Collection (ATCC). WM6026 (*lacIq rrnB3* Δ*lacZ*4787 *hsdR*514Δ*araBAD*567 Δ*rhaBAD568* rph-1 attλ::pAE12(ΔoriR6K-cat:: Frt5), *ΔendA*::Frt *uidA*(ΔMluI)::pir attHK::pJK1006(ΔoriR6K-cat::Frt5; trfA::Frt) ΔdapA::Frt) and broad host plasmid pRL814 bearing GFP and spectinomycin resistance gene (*aadA1*) were from Dr. Patricia Kiley and Dr. Robert Landick (University of Wisconsin, Madison). pRL814 was used as the cloning vector to generate complementation plasmids.

ZM4 deletion mutants were constructed as described before (18) using oligonucleotides listed in Table 1. Briefly, chromosomal 500 bp regions directly upstream and downstream of a target gene were amplified from ZM4 using Q5 Polymerase and primers listed in Table 1. Both fragments were inserted into a SpeI site of non-replicating plasmid pPK15543 (GFP, CmR) by Gibson assembly (18). The reaction mixture was transformed into chemically competent Mach 1. Sequences of the plasmids, isolated from chloramphenicol resistant Mach 1 transformants, were confirmed by Sanger sequencing. pPK15543 bearing sequences flanking a target gene, was introduced to ZM4 by conjugation using WM6026 (DAP-) as a donor strain as described in Felczak *et al* (21). Selection for colonies with the plasmid integrated into chromosome was for Cm resistance and DAP-independence. One colony of a ZM4-CmR conjugant was used to inoculate ZRMG without the antibiotic and culture was grown for at least 10 generations. 10^4^ CFU were spread on 100 ZRMG plates without chloramphenicol and left for 48 hours to grow. Colonies were screened for loss of GFP fluorescence using a blue light illuminator. Conjugation to ZM4 and resolution of primary integrants to obtain ZM4Δ*ndh* by recombination was performed in aerobic conditions. To obtain ZM4Δ*cydAB* and Δ*cydA*BΔ*ndh*, conjugation and the following steps were performed in anaerobic chamber. In this case, plates containing single colonies were left in 4°C outside of the chamber for 24 hours for maturation of fluorescence before screening for non-fluorescent colonies. Colony PCR from primers flanking a target gene was used to confirm the gene deletion in non-fluorescent colonies. The double mutant, Δ*cydA*BΔ*ndh*, was constructed by conjugating pPK15543 bearing regions flanking *ndh* to ZM4Δ*cydA*B strain followed by selection as for single knockout strain.

**Table 1.**
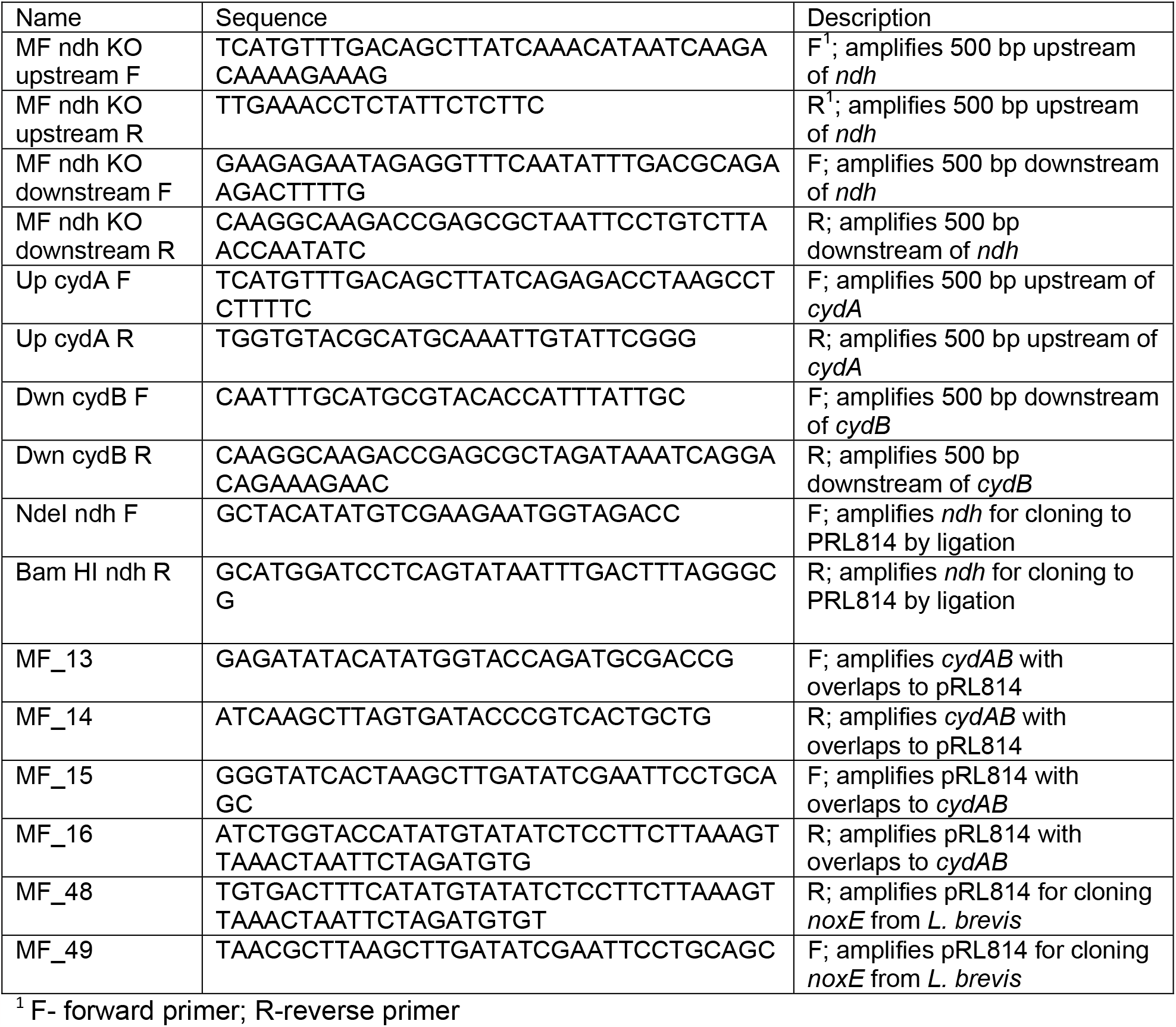
Oligonucleotides

Plasmids pRL*ndh*, pRL*cydAB*, pRL*noxE*, were constructed by introducing the indicated gene into pRL814 to replace *gfp*. All primers used in cloning are listed in Table 1. Specifically, *ndh* (ZMO1113) was amplified from ZM4 genomic DNA using primers which added NdeI and BamHI restriction sites at 5’ and 3’ ends, respectively. The fragment was then digested with above enzymes and introduced to NdeI / BamHI digested pRL814 by ligation. *cydAB* (ZMO1571-1572) was amplified from ZM4 gDNA using primers MF_13 and MF_14 and introduced to pRL814 amplified with primers MF_15 and MF_16 by Gibson assembly. A gBlock (IDTDNA) was used for cloning of *noxE* from *Lactobacillus brevis* (AF536177.1) to pRL814. GGAGATATACAT and GCTTGATATCGA overhangs were added at the 5’ and 3’ terminus, respectively, of *noxE* gene sequence. The fragment was cloned by Gibson assembly to pRL814 amplified using primers MF_48 and MF_49. All reaction mixtures were routinely transformed to Mach 1 and colony PCR was used to confirm insertion of the gene into pRL814. Finally recombined pRL814 plasmids were sequenced by the Sanger method to confirm the correctness of the construct.

### DNA Extraction and Sequencing

Genomic DNA from ZM4Δ*cydAB* isolant was extracted with DNeasy UltraClean Microbial Kit (Qiagen) and was further prepared for sequencing using NEBNext Ultra FS II kit (NEB) with the miniaturized protocol (22). Prepared NGS libraries were sequenced using Illumina NovaSeq 6000. Sample sequences were compared to reference ZM4 (ATCC31821) NCBI reference (NZ_CP023715.1) with breseq (23) using analysis pipeline available at https://github.com/mikewolfe/Bacterial_reseq.

### Bacterial growth

Strains were grown in indicated media in an anaerobic chamber overnight. Spectinomycin was added when indicated. In the morning, cultures were diluted to OD_600_=0.1 in 5 ml of the same media. For aerobic growth bacteria were grown in glass tubes covered loosely with plastic caps. For anaerobic growth, cultures were grown in Hungate tubes. In this case, after dilution of overnight cultures with fresh, anaerobic media, the tubes were closed with rubber stoppers and secured with aluminum crimps in anaerobic chamber. Aerobic and anaerobic cultures were then incubated with shaking at 275 rpm at 30°C outside of the chamber. Samples were taken periodically for cell density and HPLC analysis.

### Complementation of Δ*cydAB* growth from pRL*noxE*

Growth complementation with pRL*noxE* was performed in Zymomonas Rich Defined Medium (ZRDM) because we observed growth inhibition of ZM4 after expression of pRL*noxE* in the standard ZRMG medium. Δ*cydAB* or ZM4 bearing the vector or the complementing plasmid were grown in ZRDM supplemented with spectinomycin, overnight, in anaerobic chamber. In the morning, cultures were diluted to OD_600_=0.1 in fresh aerobic ZRDM with spectinomycin. IPTG was added to final concentrations as indicated. Wells of 96-well, clear, flat bottom microplates (Greiner) were filled with 200 µl of culture in triplicate. Bacteria were grown aerobically in 30°C incubator with shaking at speed 500 using a portable shaker (IKA MS 3 digital) for 48 hours. Absorbance at 600 nm was measured periodically by plate reader (Synergy H1 BioTek). At the end of the experiment, cultures from technical replicates were combined and OD_600_ was measured by a spectrophotometer with a 1-cm pathlength.

### HPLC analysis

Samples were taken at times indicated and stored in -20°C until used. Just before HPLC analysis, samples were thawed, centrifuged at maximum speed in benchtop microcentrifuge for 10 minutes and clear supernatants were used for analysis. HPLC chromatography was performed on Aminex HPX-87H, 300 × 7.8 mm column, at 50°C. Metabolites were eluted with 5 mM sulfuric acid at 0.6 ml/minute and detection was by refraction index (RI) based on the standards run alongside.

### Oxygen uptake measurement

Bacteria were grown overnight in ZRMG or ZRDM at 30°C in anaerobic chamber. Cultures were diluted in fresh, aerobic medium to initial OD_600_ indicated in experiments. The dilution factor was determined experimentally to adjust for O_2_ uptake rate characteristic for each strain that allowed for near 100% air saturation after 30 minutes of shaking in a plate reader. Wells of 96-well, flat bottom OxoPlate (OP96C, PrecisionSensing) were filled with 200 µl of each culture in triplicate. Control wells were filled with 400 µl of oxygen-free calibration solution (Cal 0%) or 200 µl of 100% air-saturated calibration solution (Cal 100%), in quadruplicate, as recommended by the manufacturer. The wells containing Cal 0% were covered with adhesive foil. Cal 0% was prepared by dissolving 0.15 g of Na_2_SO_3_ in 15 ml water in 15 ml, tightly closed VWR vial. Cal 100% was obtained by vigorous vortexing of 20 ml of H_2_O in 50 ml VWR conical tube for two minutes and subsequent gentle moving of liquid for one minute to reduce air oversaturation. The vial was closed and both standards were used immediately. The plate was covered with the lid and allowed to shake in a plate reader in aerobic conditions for 30 minutes at 30°C to stabilize the temperature and to allow for expression of oxygen-inducible genes. After this time, shaking was stopped and reference, (540/590) and indicator (540/650) fluorescence, were measured every 3.2 minutes for one hour. Oxygen partial pressure as % of air saturation (pO_2_), was calculated from the formula: pO_2_ = 100*(k0/IR -1)/(k0/k100-1) where IR = Fluorescence indicator/Fluorescence reference (Fi/Fr) of unknown sample; k0 = Fi/Fr of Cal 0% and; k100 = Fi/Fr of Cal 100%. Final value of O_2_ concentration in mg/l was obtained by multiplying pO_2_ by conversion factor of 0.091. O_2_ consumption rate was calculated from the linear part of the slope and normalized to OD_600_=1.

## Acknowledgements

This material is based upon work supported by the Great Lakes Bioenergy Research Center, U.S. Department of Energy, Office of Science, Office of Biological and Environmental Research under Award Number DE-SC0018409. We thank Bailey Marshall for conducting the genome re-sequencing, and Dr. Mike Wolfe for helpful discussions regarding *Z. mobilis* regulons.

## References

1. Flamholz A, Noor E, Bar-Even A, Liebermeister W, Milo R. 2013. Glycolytic strategy as a tradeoff between energy yield and protein cost. Proc Natl Acad Sci U S A 110:10039–10044.

2. Brenac L, Baidoo EEK, Keasling JD, Budin I. 2019. Distinct functional roles for hopanoid composition in the chemical tolerance of Zymomonas mobilis. Mol Microbiol 112:1564–1575.

3. Fuchino K, Wasser D, Soppa J. 2021. Genome copy number quantification revealed that the ethanologenic alpha-proteobacterium Zymomonas mobilis is polyploid. Front Microbiol 12:705895.

4. Bringer S, Finn RK, Sahm H. 1984. Effect of oxygen on the metabolism of Zymomonas mobilis. Arch Microbiol 139:376–381.

5. Belauich JP, Senez JC. 1965. Influence of aeration and of pantothenate on growth yields of Zymomonas mobilis. J Bacteriol 89:1195–1200.

6. Kalnenieks U, Galinina N, Toma MM, Poole RK. 2000. Cyanide inhibits respiration yet stimulates aerobic growth of Zymomonas mobilis. Microbiology 146:1259–66.

7. Kalnenieks U, de Graaf AA, Bringer-Meyer S, Sahm H. 1993. Oxidative phosphorylation in Zymomonas mobilis. Arch Microbiol 160:74–79.

8. Felczak M, TerAvest M. 2022. Zymomonas mobilis ZM4 utilizes an NADP+-dependent acetaldehyde dehydrogenase to produce acetate. J Bacteriol 204:e00563–21.

9. Jones-Burrage SE, Kremer TA, McKinlay JB. 2019. Cell aggregation and aerobic respiration are important for Zymomonas mobilis ZM4 survival in an aerobic minimal medium. Appl Environ Microbiol 85:e00193–19.

10. Hayashi T, Furuta Y, Furukawa K. 2011. Respiration-deficient mutants of Zymomonas mobilis show improved growth and ethanol fermentation under aerobic and high temperature conditions. J Biosci Bioeng 111:414–419.

11. Kalnenieks U, Galinina N, Strazdina I, Kravale Z, Pickford JL, Rutkis R, Poole RK. 2008. NADH dehydrogenase deficiency results in low respiration rate and improved aerobic growth of Zymomonas mobilis. Microbiology 154:989–994.

12. Rutkis R, Strazdina I, Balodite E, Lasa Z, Galinina N, Kalnenieks U. 2016. The Low Energy-Coupling Respiration in Zymomonas mobilis Accelerates Flux in the Entner-Doudoroff Pathway. PLoS One 11:e0153866.

13. Strohdeicher M, Neuß B, Bringer-Meyer S, Sahm H. 1990. Electron transport chain of Zymomonas mobilis. Arch Microbiol 154:536–543.

14. Strazdina I, Balodite E, Lasa Z, Rutkis R, Galinina N, Kalnenieks U. 2018. Aerobic catabolism and respiratory lactate bypass in Ndh-negative Zymomonas mobilis. Metab Eng Commun 7:e00081.

15. Sootsuwan K, Lertwattanasakul N, Thanonkeo P, Matsushita K, Yamada M. 2008. Analysis of the respiratory chain in ethanologenic Zymomonas mobilis with a cyanide-resistant bd-type ubiquinol oxidase as the only terminal oxidase and its possible physiological roles. J Mol Microbiol Biotechnol 14:163–75.

16. Balodite E, Strazdina I, Galinina N, McLean S, Rutkis R, Poole RK, Kalnenieks U. 2014. Structure of the Zymomonas mobilis respiratory chain: oxygen affinity of electron transport and the role of cytochrome c peroxidase. Microbiology 160:2045–52.

17. Strazdina I, Kravale Z, Galinina N, Rutkis R, Poole RK, Kalnenieks U. 2012. Electron transport and oxidative stress in Zymomonas mobilis respiratory mutants. Arch Microbiol 194:461–71.

18. Lal PB, Wells FM, Lyu Y, Ghosh IN, Landick R, Kiley PJ. 2019. A markerless method for genome engineering in Zymomonas mobilis ZM4. Front Microbiol.

19. Felczak MM, Jacobson TB, Ong WK, Amador-Noguez D, TerAvest MA. 2019. Expression of Phosphofructokinase Is Not Sufficient to Enable Embden-Meyerhof-Parnas Glycolysis in Zymomonas mobilis ZM4. Front Microbiol.

20. Martien JI, Hebert AS, Stevenson DM, Regner MR, Khana DB, Coon JJ, Amador-Noguez D. 2019. Systems-level analysis of oxygen exposure in Zymomonas mobilis: implications for isoprenoid production. mSystems 4:e00284–18.

21. Felczak MM, Bowers RM, Woyke T, TerAvest MA. 2021. Zymomonas diversity and potential for biofuel production. Biotechnol Biofuels 14:112.

22. Li H, Wu K, Ruan C, Pan J, Wang Y, Long H. 2019. Cost-reduction strategies in massive genomics experiments. Mar Life Sci Technol 1:15–21.

23. Deatherage DE, Barrick JE. 2014. Identification of mutations in laboratory-evolved microbes from next-generation sequencing data using breseq. Methods Mol Biol 1151:165–188.

